# Genome-wide association analysis of flowering date in a collection of cultivated olive tree

**DOI:** 10.1101/2024.06.10.598200

**Authors:** Laila Aqbouch, Omar Abou-Saaid, Gautier Sarah, Lison Zunino, Vincent Segura, Pierre Mournet, Florelle Bonal, Hayat Zaher, Ahmed El Bakkali, Philippe Cubry, Evelyne Costes, Bouchaib Khadari

## Abstract

Flowering date in perennial fruit trees is an important trait for fruit production. Depending on the winter and spring temperatures, flowering of olive may be advanced, delayed, or even suppressed. Deciphering the genetic control of flowering date is thus key to help selecting cultivars better adapted to the current climate context. Here, we investigated the genetic determinism of full flowering date stage in cultivated olive based on capture sequencing data of 318 genotypes from the worldwide olive germplasm bank of Marrakech, Morocco. The genetic structure of this collection was organized in three clusters that were broadly attributed to eastern, central, and western Mediterranean regions, based on the presumed origin of genotypes. Flowering dates, collected over seven years, were used to estimate the genotypic best linear unbiased predictors, which were then analyzed in a genome-wide association study. Loci with small effects were significantly associated with the studied trait, by either a single- or a multi-locus approach. The three most robust loci were located on chromosomes 01 and 04, and on a scaffold, and explained 7.1%, 6.2%, and 6.5 % of the trait variance, respectively. A significantly higher accuracy in the best linear unbiased predictors of flowering date prediction was reported with Ridge-compared to LASSO-based genomic prediction model. Along with genomic association results, this suggests a complex polygenic determinism of flowering date, as seen in many other fruit perennials. These results and the screening of associated regions for candidate genes open perspectives for further studies and breeding programs targeting flowering date.

## 2. Introduction

Flowering date in fruit perennial trees is known to be influenced by temperature, specifically during periods of accumulation of chill and heat requirements (Guo et al., 2014). Increasing temperatures during winter can result in difficulties in chilling requirements fulfillment and may delay flowering date (Atkinson et al., 2013). In contrast, the increase in temperatures during spring advances the flowering date (Grab and Craparo 2011). This can increase frost damage risk (Saxe et al., 2001), and result in several morphological disorders, such as bud burst delay, low burst rate, irregular floral or leaf budbreak and poor fruit set (Dirlewanger et al., 2012). In allogamous species with a self-incompatibility reproductive system, it can also cause asynchrony between compatible varieties (Dirlewanger et al., 2012). This may disturb pollination and consequently, fruit production (Atkinson et al., 2013).

Flowering date has been shown to be quantitatively inherited in fruit trees, several Quantitative Trait Loci (QTL) have been detected in bi- or multi-parental populations of apple tree (Allard et al., 2016), peach (Li et al., 2023), and apricot (Kitamura et al., 2018). More recently, Genome-Wide Association Study (GWAS) have been conducted on several fruit tree species (e.g. Watson et al., 2024). However, no similar study has been conducted so far on the cultivated olive tree, an emblematic species of the Mediterranean Basin (MB), despite the region being known to be particularly affected by the current global warming (Pardo et al., 2023).

GWAS is one of the methods used to discover genetic variations affecting complex traits (Abdellaoui et al., 2023). Unlike QTL mapping studies, GWAS can investigate associations within populations where relatedness among individuals is variable, and even when the relatedness is unknown (Atwell et al., 2010). To handle spurious associations, several factors have to be considered, including population structure and linkage disequilibrium (LD), which could associate non-causal variants in LD with the causal variants to the trait (Uffelmann et al., 2021).

The olive tree (*Olea europaea* L.) is often considered as an iconic species of MB. It is believed that olive has been domesticated around 6000 years ago, with a main domestication event in the eastern MB supported by several studies (Khadari and El Bakkali, 2018). It remains unclear whether subsequent diversification followed the first domestication (Khadari and El Bakkali, 2018), or if a second independent domestication event occurred in the central Mediterranean area (Diez et al., 2015). The cultivated olive tree is diploid, and 23 chromosomes have been assembled (Julca et al., 2020). Four assembled genomes are currently available for the species *Olea europaea var. europaea*: two versions of cv. *Farga*: Oe6 version (Cruz et al., 2016) and Oe9 version (Julca et al., 2020), cv. *Picual* (Jiménez-Ruiz et al., 2020) and cv. *Arbequina* (Rao et al., 2021). The last version of *Farga* estimated the length of the olive genome to be approximately 1.3 Gb, with 7.3 Mb corresponding to scaffolds and 54 Kb to contigs (Julca et al., 2020). This genome was the last one available when we started the present study. A more recent assembly of the *Arbequina* cultivar was published afterwards that has estimated a similar genome length with 1.25 Gb on chromosomes (Rao et al., 2021).

Several germplasm collections of olive trees have been constituted, the two most extensive being the Worldwide Olive Germplasm Bank of Marrakech, Morocco (WOGBM) and Cordoba, Spain (WOGBC) (El Bakkali et al., 2019). The genetic structure of the WOGBM has been investigated using Simple Sequence Repeat (SSR) markers (El Bakkali et al., 2019), while that of the WOGBC relies on SSR (Diez et al., 2015) and Expressed Sequence Tag Single Nucleotide Polymorphism (EST-SNP) markers (Belaj et al., 2022). These analyses resulted in the detection of three distinct genetic clusters, corresponding to the assumed geographical areas of origin of cultivars, with a large proportion of non-assigned individuals.

Those collections have been phenotyped for several traits, in particular, flowering date. A large variation in this trait between years has been observed in the WOGBM (Abou-Saaid et al., 2022). As other fruit tree species, this variability is assumed to rely on temperature sensing during winter and spring (Guo et al., 2014). In addition, the olive tree presents the particularity to require low temperature for floral induction (Haberman et al., 2017). Therefore, in olive tree, winter temperatures not only impact the flowering dates but also its occurrence (Benlloch-González et al., 2018). Under the current climate change situation that deeply modifies temperature regimes, the major risk for olive trees concerns the synchrony between compatible varieties, which may disturb their cross-pollination. Indeed, the sexual reproductive system of olive is allogamous due to a self-incompatibility system (Saumitou-Laprade et al, 2017). Since successful pollination is a main factor in fruit development, flowering date is a key trait for the success of the olive tree reproductive cycle, upon which the uniformity and quality of fruit production depend (El Yaacoubi et al., 2014).

The main purpose of our study was to explore the genetic determinism of flowering date in cultivated olive, based on a specific phenological stage, the full flowering date (FFD). For this intent, the large panel of genetic diversity from the WOGBM and a new high-quality SNP data that we developed through capture sequencing were used in a GWAS. This new genotypic data was first validated through a genetic structure analysis before considering it for the GWAS.

## 3. Results

### Characterization and distribution of SNPs in the cultivated olive genome

We initially sequenced 335 genomic libraries. The raw sequencing data ranges from 1,590 read pairs for the *Atounsi Setif* (MAR00516) genotype to 39,801,319 read pairs for the *Aggezi Shami* (MAR00480) genotype, with a mean of 8,603,434 read pairs (Figure S1). The *Aharoun* (MAR00447) genotype was filtered out (quality reads below 30). After cleaning, the read pairs count ranged from 1,514 to 39,231,314, with a mean of 8,488,947 (Figure S1).

We mapped our reads to the latest version of the Farga Oe9 reference genome assembly (Julca et al., 2020). A mean of 98.82 % of the reads were mapped on the Farga genome and tagged as properly paired. The mapping rate ranged from 84.68% (*Atounsi Setif*) to 99.59% (*Sayali* (MAR00287)). The genotype *Azeradj Tamokra* (MAR00448) was removed (mapping rate of 0%). The mean enrichment rate in targeted sequences was 39 times (Table S1).

A total of 64,835,479 variants were initially identified among 333 samples (*Azeradj Tamokra* and *Aharoun* were filtered out). After removing experimental duplicates, biological replicates, and individuals whose genomic libraries were not captured, 325 unique genotypes remained (Table S2). After handling filtration steps to ensure retrieving SNP of high-quality, we retained 235,825 SNPs across 318 genotypes (Table S2). These SNPs were then used for genetic structure and PCA analyses. Additional filters (retaining only nuclear markers, filtering on Minor Allele Frequency, and imputation of missing data) resulted in 118,948 SNPs across 318 genotypes, which were used for GWAS and genomic prediction analyses (Table S2). Of these SNPs, 49.2% were in the targeted region by the baits, while the remaining were in the non-target region. Approximately 50% of the filtered SNPs were located on chromosomal regions, while the rest of SNPs were found on scaffolds.

### Three genetic clusters are identified in the WOGBM collection

The sNMF approach (Frichot et al., 2014) was used to analyze population structure using 235,825 high-quality SNPs from 318 genotypes. The sNMF approach estimated individual ancestry coefficients and helped determine the number of ancestral populations (Table S3). We set the number of clusters to three based on the cross-entropy criterion (Figure S2).

A genotype was assigned to a genetic cluster if it had a minimum of 70% ancestry estimation within that cluster. Genotypes not reaching a 70% assignment to any of the three genetic clusters were classified as non-assigned. Out of the 318 genotypes, 79 were assigned to the ancestry cluster K1 (from 71% to 100%). This group of genotypes was denoted C1 in the following. 33 genotypes were assigned to the ancestry cluster K2 (from 71% to 100%). This group of genotypes was denoted C2. 71 genotypes were assigned to the ancestry cluster K3 (from 72% to 100%). This group of genotypes was denoted C3. The remaining 135 genotypes were non-assigned and their group was denoted as the M group (Figure 1). A PCA performed using the same genotypic dataset highlighted that the genotypes from the three genetic groups, C1, C2, and C3, were clearly separated on the plot of the first two components (Figure 2). The first principal component accounted for 9.5% of the genetic variability and separated C2 from C1 and C3. The second principal component accounted for 5% of the genetic variability and separated C1 from C2 (Figure S3, Figure 2). PC3 explained 2.6% of the genetic variability (Figure S3). The genotypes in the C3 group appeared to be more closely related compared to those assigned to the other two groups, C1 and C2, whether on the PC1-PC2 plot (Figure 2) or the PC2-PC3 plot (Figure S4). Non-assigned individuals were widely spread in the region between the three groups on PC1 and PC2 (Figure 2).

**Figure 1.**
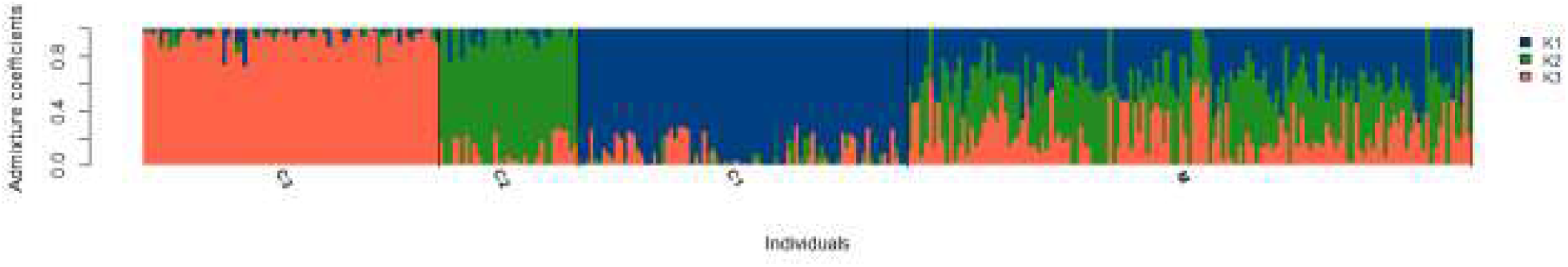
Admixture coefficients as inferred by sNMF analysis (Frichot et al., 2014) for the 318 genotypes of Worldwide Olive Germplasm Banks of Marrakech (WOGBM) using 235,825 SNPs. Bars are ordered by assignment to genetic clusters K1, K2, or K3. Groups of genotypes were named C1 for those assigned to the genetic cluster K1, C2 for those assigned to the genetic cluster K2, C3 for those assigned to the genetic cluster K3, and M for genotypes non-assigned to a genetic cluster.

**Figure 2.**
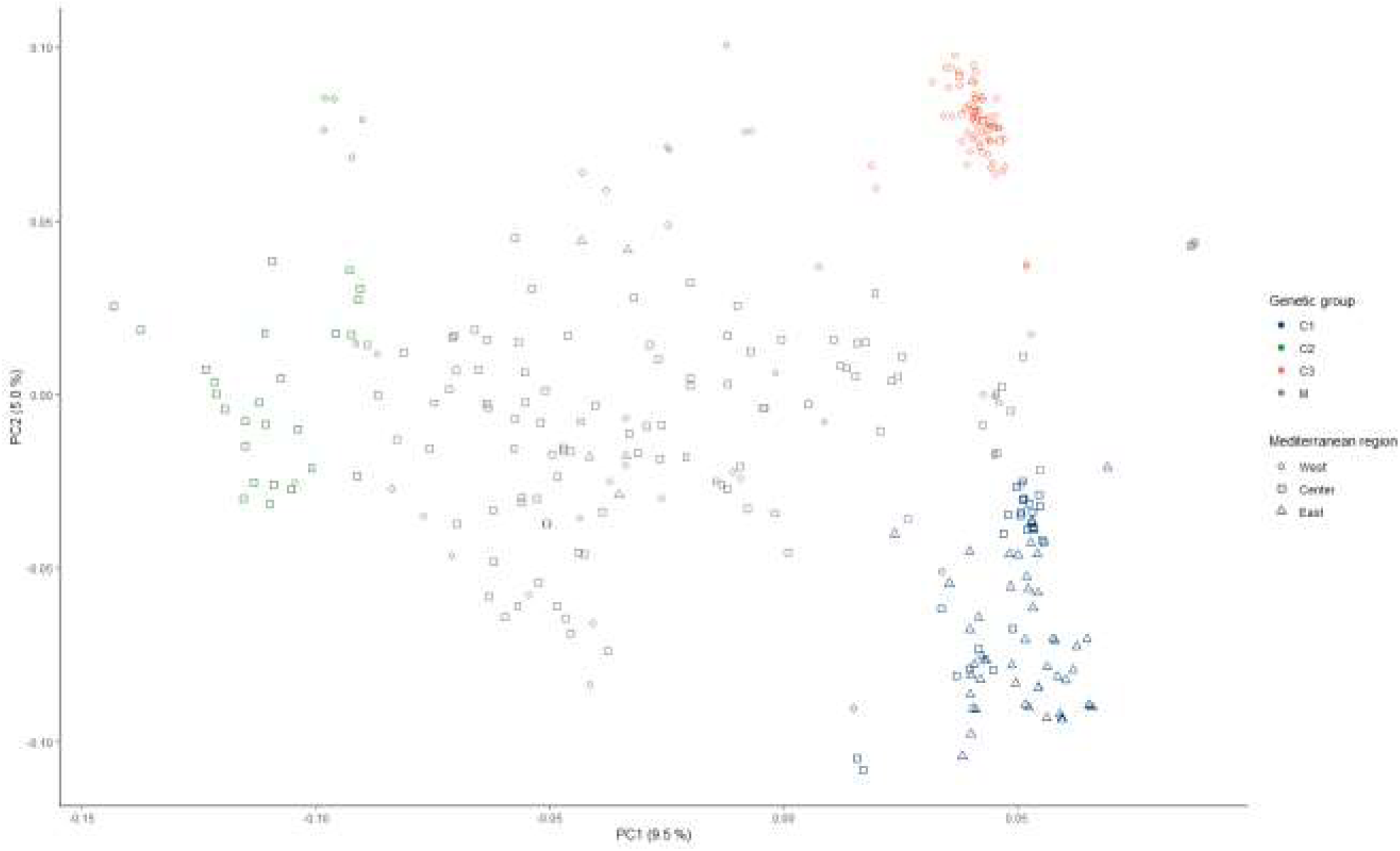
Projection of the 318 genotypes from WOGBM on the first two principal components (PC) of a PC analysis based on 235,825 SNPs. Colors blue, green, and orange indicate the group to which each genotype was assigned (C1, C2, C3), and grey indicates the non-assigned genotypes (M). Circles, squares, and triangles indicate genotypes that are assumed to originate from the western, central, and eastern regions of the Mediterranean basin (MB), respectively. The east corresponds to Cyprus, Egypt, Greece, Lebanon, and Syria; the center corresponds to Algeria, Croatia, France, Italy, Slovenia, and Tunisia; and the west corresponds to Algeria, Croatia, France, Italy, Slovenia, and Tunisia.

The information regarding the assumed origin of genotypes in the WOGBM (El Bakkali et al., 2019) was crossed with the genetic structure analysis results. We ordered the barplot displaying individually estimated ancestries of genotypes based on the assumed geographical origin. We started ordering from the western Mediterranean on the left and progressing towards the eastern Mediterranean on the right according to the country of origin indicated in their passport data (Figure 3). This representation suggests a geographical basis for the genetic structure. To further explore this geographically based genetic structure hypothesis, we confronted information about the genotype’s genetic cluster assignment, following the criteria presented above (i.e. an individual is assigned to a cluster if they have a minimum of 70% ancestry estimation within that cluster), with information about the supposed country of origin (Table S4). 70% of genotypes of the C1 group had a supposed origin from Cyprus, Egypt, Greece, Lebanon and Syria (eastern MB). 79% of the C2 group genotypes were indicated in their passport data as originating from Algeria, Croatia, France, Italy, Slovenia and Tunisia (central MB). 93% of the C3 group genotypes were supposed to originate from Morocco, Spain, and Portugal (western MB). The non-assigned group of genotypes consists of 70% of genotypes supposed to originate from the central MB (Table S4).

**Figure 3.**
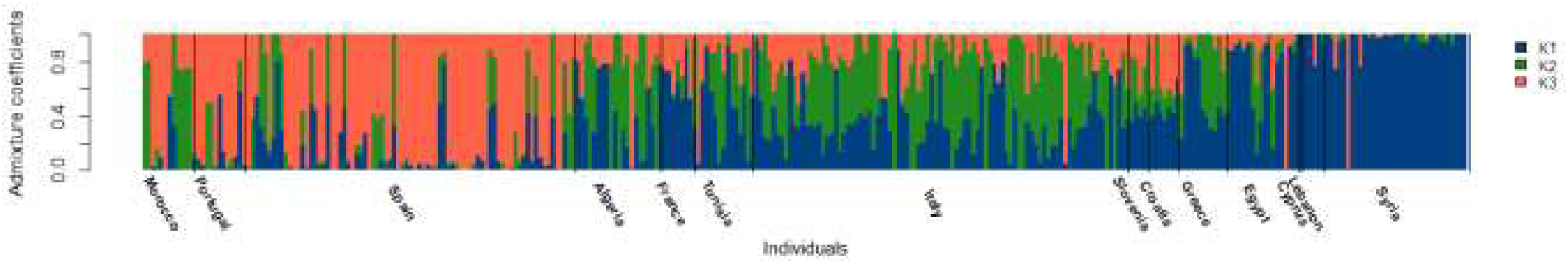
Admixture coefficients as inferred by sNMF analysis (Frichot et al., 2014) of the 318 genotypes of WOGBM using 235,825 SNPs. Bars indicate the proportion of assignment to genetic clusters K1, K2, or K3 and are sorted by the assumed geographic origin of genotypes, from western to eastern Mediterranean regions.

### Flowering date is different among genetic groups

The Best Linear Unbiased Predictor (BLUP) of the genotype effect was estimated using a mixed model that included genotype, year, and the interaction between genotype and year effects based on data of seven years. The collection contained at least three trees for each genotype. The variance of the phenotypes, based on raw data, was 98.77 calendar days. After the mixed model estimation, the variance attributed to the genotypic effect was 4.12 days, the variance of the interaction between genotypes and years was 4.61 days, whereas the residual variance was 5.53 days. Based on the variance components issued from the model, the broad-sense heritability was estimated at 0.84, indicating a relatively high value. The genetic BLUP of flowering date in the whole collection (331 genotypes) follows a normal distribution (Shapiro-Wilk, p-value = 0.97), with a mean value of 116.37 calendar days. The range spans 10.4 days, with minimum and maximum values of 110.8 days for the genotype *Borriolenca* and 121.1 days for the genotype *Ogliarola del Bradano* respectively (Figure S5). The distribution of the genetic BLUP of flowering dates was compared across the different genetic groups C1, C2, and C3 (Figure 4). A significant difference in the distribution of genotypic BLUP of FFD was observed among genetic clusters based on a Mann-Whitney pairwise comparison test (Table S5). C1 genotypes exhibited the earliest FFD values, with a mean of 115.47 calendar days, including genotypes such as *Karme* and *Minekiri*. C2 genotypes flowered the latest, with a mean value of 117.55 days, including genotypes such as *Ogliarola del Bradano* and *Olivastra di Populonia*. C3 exhibits an intermediate flowering date compared to C1 and C2, with a mean value of 116.53 days, including genotypes such as *Negrillo de Iznalloz* and *Manzanilla de Agua*. C1 genotypes were highly distinct from both C2 and C3 ones, according to the p-values of the Mann-Whitney test (Table S5).

**Figure 4.**
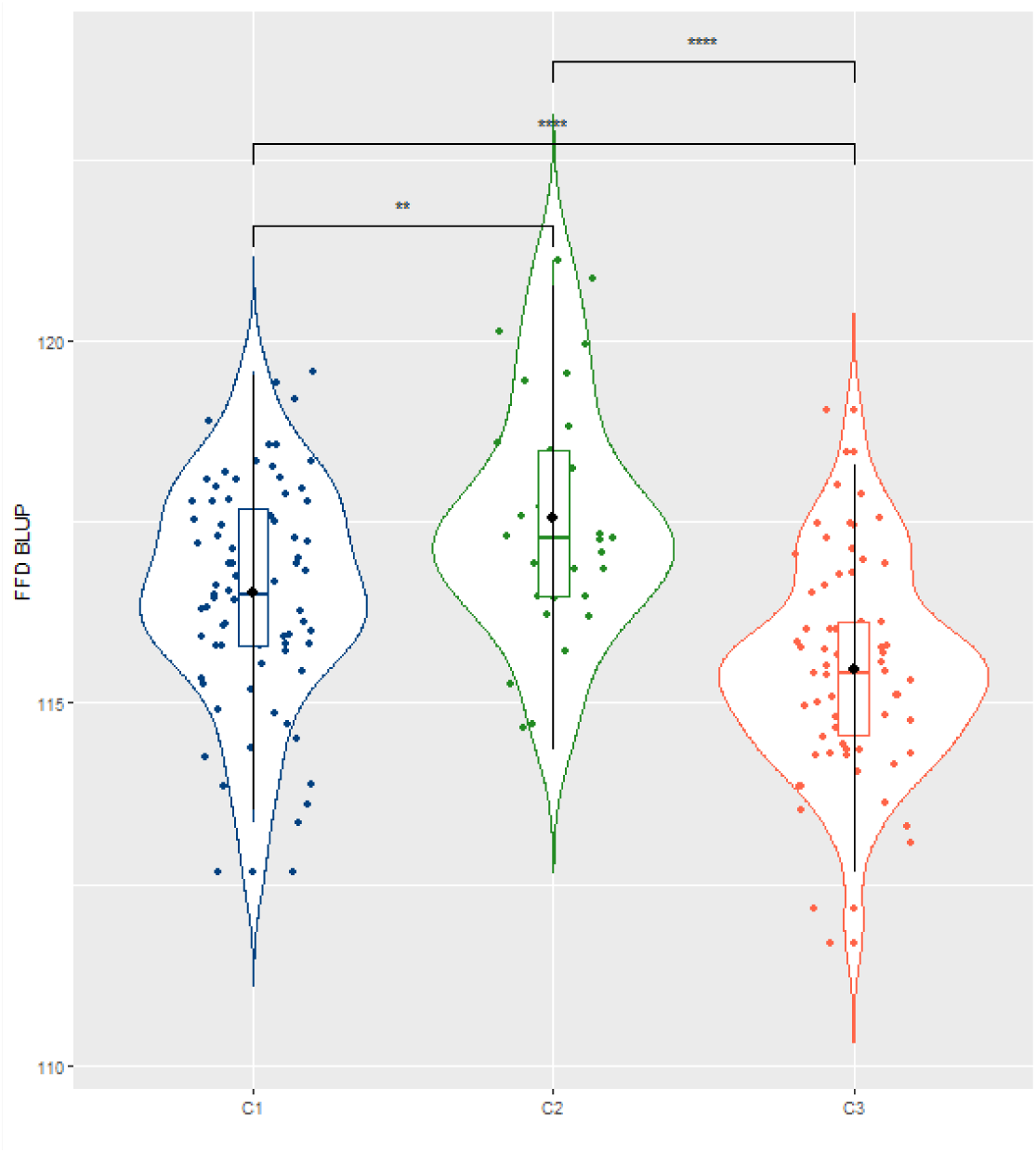
Distribution of the genetic BLUP of FFD depending on the genetic groups (C1 in blue, C2 in green, and C3 in orange) with pairwise significance of their difference according to the Wilcoxon-Mann-Whitney test (Wilcoxon, 1945). Levels of significance: ns (not significant); * (p<0.05); ** (p<0.01); *** (p<0.001). Black circles indicate the mean value, the horizontal bar the median value, and the box plot the first and third quartile of each distribution, respectively.

### Three genomic regions are associated with FFD using single-locus and multi-locus association analyses

Before performing the association study, we tested three linear mixed models that account for structure and/or kinship effects. The structure was considered as a fixed effect (as assessed by the ancestry matrix obtained from the sNMF run that exhibited the lowest cross-entropy value at the considered K, Q model) while the kinship was considered as the covariance matrix of a random effect separately (u model) or jointly (u+Q model). We tested two kinship matrices: Weir & Goudet (Weir and Goudet, 2017), recommended for populations with related individuals (Goudet et al., 2018), and VanRaden Kinship (VanRaden, 2008), widely used in association studies. We found that the best model was the one considering kinship only, regardless of the considered kinship matrix (Table S6). This model (u model) was thus retained to investigate the genetic determinism of the FFD trait using a GWAS approach. We firstly used a single-locus mixed-model approach, implemented in the R package MM4LMM (Laporte et al., 2022), and complemented it with a multi-locus method, MLMM (Segura et al., 2012). The two distinct kinship matrices (Weir & Goudet and VanRaden) previously described were tested for each of the two approaches, resulting in four analyses.

Associations were tested between the genotypic BLUP of FFD (Table S7) and 118,948 high-quality SNP datasets obtained after applying all filtering criteria (Table S2) from 318 genotypes in the WOGBM collection. The empirical significance threshold for MM4LMM was set at a 5% FDR, a commonly used criterion (Nelson et al., 2017). For MLMM, the significance threshold was set at 9.6E-6, which corresponds to the p-value of the least significant SNP in the initial run analysis of MM4LMM using the Weir & Goudet kinship (Weir and Goudet, 2017).

The single-locus approach resulted in 23 significantly associated SNPs when using the Weir & Goudet kinship (Figure 5 A, Figure 5 B, Table S8), while no SNP was detected when using the VanRaden kinship (Table S8). P-values of the significant SNPs ranged from 1.5E-07 for the “Oe9_LG01_9017771” SNP to 9.6E-06 for the “Oe9_LG05_12679503” SNP (Table S8).

**Figure 5.**
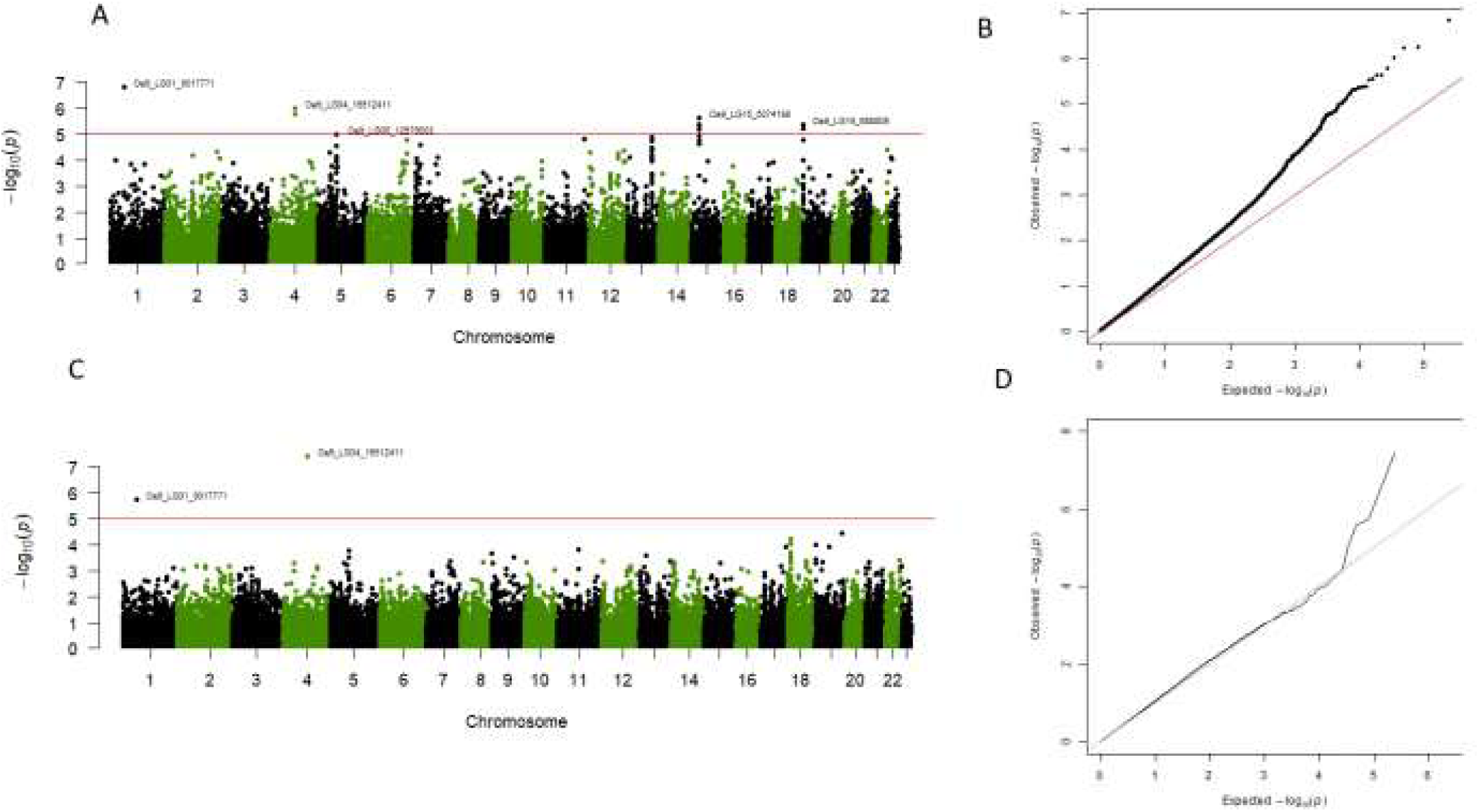
Manhattan plot of the GWAS study of genotypic BLUP of FFD using Weir & Goudet kinship (only chromosomal regions are shown in the plot). A. Manhattan plot based on the single-locus approach MM4LMM. B. Q–Q plot corresponding to the MM4LMM model. C. Manhattan plot based on the multi-locus approach MLMM. D. Q–Q plot corresponding to the MLMM model. The horizontal red line in the Manhattan plots indicates the p-value that corresponds to a threshold of 5% false discovery rate (FDR) in the MM4LMM model using the Weir & Goudet kinship.

The multi-locus approach yielded six significant SNPs, depending on the kinship matrix considered. Four of them were detected using Weir & Goudet kinship, having p-values ranging from 3.74E-08 for “Oe9_LG04_16512411” SNP to 9.11E-06 for “Oe9_s06150_161951” SNP (Figure 5 C, Figure 5 D, Table S8). Three SNPs were detected using VanRaden, with p-values ranging from 4.81E-08 for the “Oe9_s07747_163567” SNP to 6.41E-06 for the “Oe9_LG04_16512411” SNP (Table S8).

A total of 26 SNPs were significantly associated with the FFD BLUPs in at least one of the four association analyses. Two SNPs, “Oe9_LG01_9017771” and “Oe9_s04305_16459”, were detected by two of the four analyses, while only one SNP, “Oe9_LG04_16512411”, was detected by three analyses (Table S8, Figure S6 A,B, and C). These three SNPs were considered as strong candidates, with “Oe9_LG04_16512411” being the most robust. The three SNPs: “Oe9_LG01_9017771”, “Oe9_s04305_16459”, and “Oe9_LG04_16512411”, explained 7.1%, 6.5%, and 6.2% of the trait’s variance, respectively (Table 1).

**Table 1.** Characterization of the three robust SNPs significantly associated with genotypic BLUP of FFD: SNP name, chromosome or scaffold number, position in base pair, allelic composition (Ref indicates the allele of reference and ALT the alternative allele), minor allele frequency (MAF), Model (MM4LMM or MLMM), Kinship matrix (Weir & Goudet or VanRaden), p-value and portion of variance explained (R2) by each SNP.

| SNP_name | Linkage group | Position (bp) | Alleles(Ref/ALT) | MAF | Model | Kinship | P_value | R2 |
| --- | --- | --- | --- | --- | --- | --- | --- | --- |
| Oe9_LG01_9017771 | Chromosome 01 | 9017771 | T/C | 0.17 | MM4LMM | Weir & Goudet | 1.50E-07 | **0.071** |
|  |  |  |  |  | MLMM | Weir & Goudet | 1.78E-06 |  |
| Oe9_s04305_16459 | s04305 | 16459 | T/C | 0.10 | MM4LMM | Weir & Goudet | 5.77E-07 | **0.065** |
|  |  |  |  |  | MLMM | VanRaden | 1.51E-06 |  |
| Oe9_LG04_16512411 | Chromosome 04 | 16512411 | G/C | 0.06 | MM4LMM | Weir & Goudet | 1.01E-06 | **0.062** |
|  |  |  |  |  | MLMM | Weir & Goudet | 3.74E-08 |  |
|  |  |  |  |  | MLMM | VanRaden | 6.41E-06 |  |

### FFD can be predicted with high accuracy using genomic prediction approach

A limited portion of the variance in the genotypic BLUP of the FFD trait was explained by the associated SNPs from the GWAS study (6.2% to 7.1% for the 3 SNPs retained as most robust). We aimed to investigate whether genomic prediction using a larger set of SNPs could account for a larger proportion of the trait’s variance.

For this purpose, we complemented the association analyses with a modeling approach based on a genome-wide analysis, using all SNPs simultaneously. This approach made use of genomic prediction models with two complementary regression approaches, Least Absolute Shrinkage and Selection Operator (LASSO) and Ridge regression (RR), respectively. LASSO estimation relies on a limited number of major effects, whereas RR is based on many minor effects. The prediction accuracy was measured by calculating Pearson’s correlation between predicted and observed values on a cross-validation setting with 5 folds and repeated one hundred times. Overall, the prediction of the FFD trait demonstrated relatively high accuracy, whether by LASSO or RR (Figure 6). The accuracy values for the RR model ranged from 0.47 to 0.79, whereas those for the LASSO model ranged from 0.31 to 0.70. The RR model achieved a significantly higher (Wilcoxon-Mann-Whitney, p-value = 6.1e-11) mean accuracy (0.64) compared to the LASSO-based model (0.55) in predicting the trait (Figure 6).

**Figure 6.**
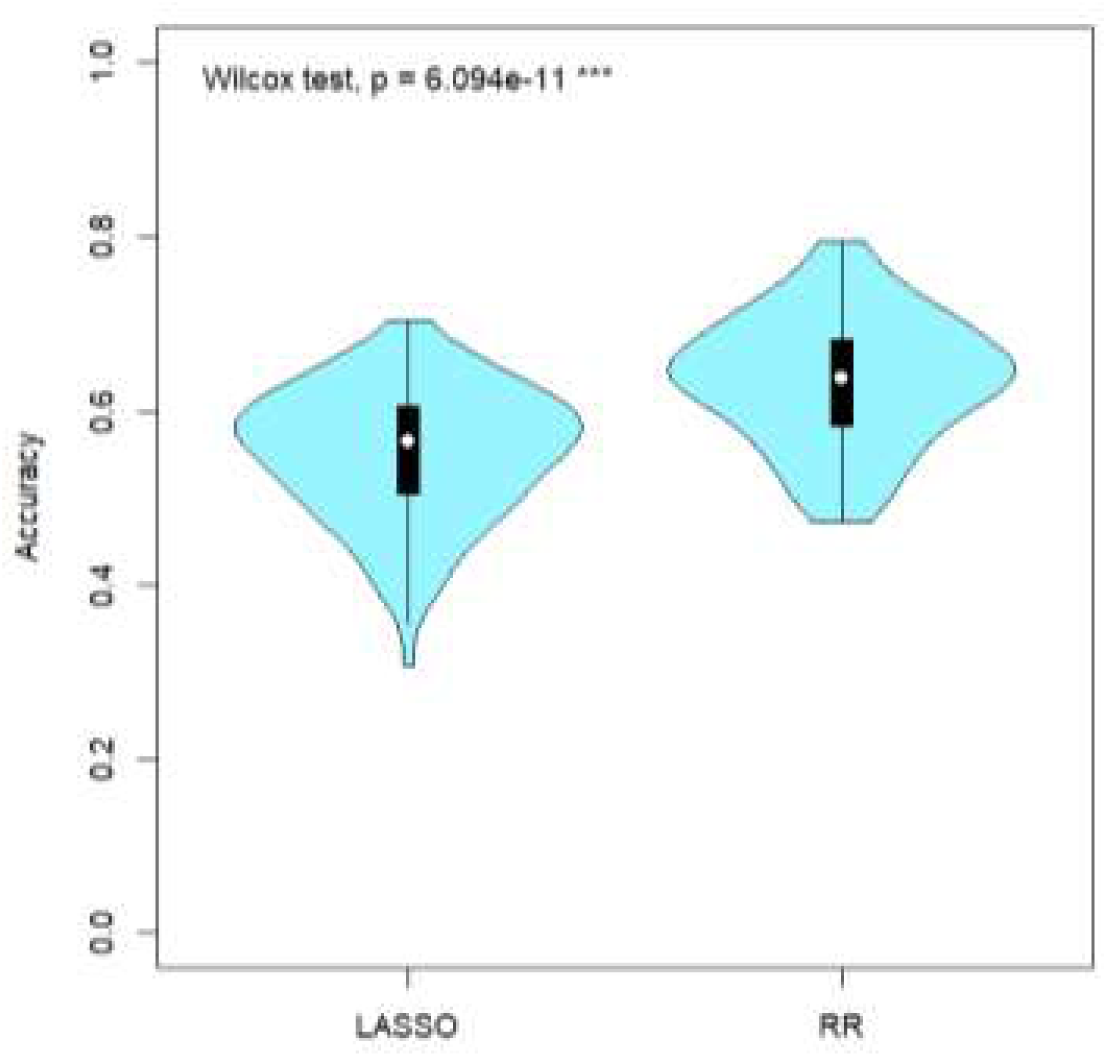
Distribution of Pearson’s correlation between predicted and observed values (accuracy) according to LASSO- and Ridge-based models based on 100 iterations. p is the p-value of the Wilcoxon-Mann-Whitney test of comparison of the two distributions (Wilcoxon, 1945). Levels of significance: ns (not significant); * (p<0.05); ** (p<0.01); *** (p<0.001). white circles indicate the mean value, and the boxplot the first and third quartile of each distribution, respectively.

### Identification of candidate genes in the genomic regions putatively associated with flowering date

We specifically examined the genomic regions neighboring the three SNPs previously identified as the most robust by single and multi-locus approaches. To ensure the inclusion of all neighboring SNPs in linkage disequilibrium (LD) in the genomic region of interest, we first analyzed the LD decay within our SNP dataset. A relatively rapid decay of LD was observed, where the average r2 values dropped within 100 bp from 0.35 which corresponds to the maximum value to 0.2 (Figure S7). Considering such a rapid LD decay, we used genomic windows of 1500 bases upstream and downstream of the associated SNP positions to retrieve candidate genes (Table 2). Based on the annotation of the reference genome (Julca et al., 2020), three genes were identified: *OE9A117378* and *OE9A084268* on Scaffold s04305 and *OE9A057547* gene on chromosome 01 (Table 2). No gene was identified within the associated genomic region on Chromosome 04 (Table 2, Table S9). We blasted the transcripts of the three genes against the UniProt database (The UniProt Consortium, 2023). A high degree of sequence similarity was identified with the *XCT* gene for the olive genes *OE9A117378* and *OE9A084268*.

**Table 2.**
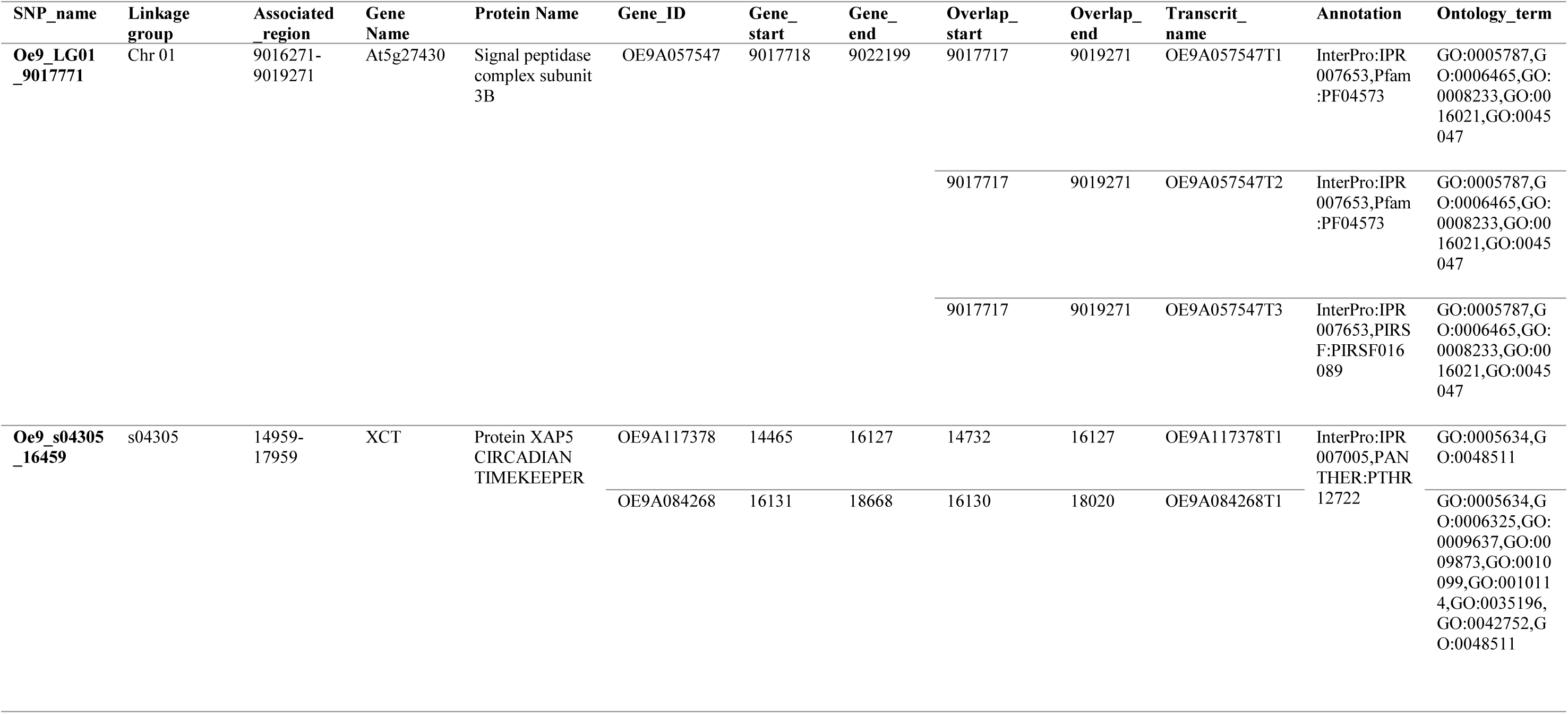
Annotation of genes found in the associated regions, corresponding to 1500pb upstream and downstream each of the three robust SNPs linked with genotypic BLUP of FFD: SNP name, chromosome (Chr) or scaffold number, interval position of the associated region from the olive reference genome Farga V2 (Julca et al., 2020); Gene and protein names based on UniProt database (The UniProt Consortium, 2023); Gene ID, position, Transcripts, respective positions indicating their overlap, annotation and ontology term from the reference genome Farga V2.

The *Oryza sativa XCT* gene exhibited 80.1% identity with the olive gene *OE9A117378*, while the *Arabidopsis thaliana XCT* gene shared 94.8% identity with the olive gene *OE9A084268*. The *XCT* gene encodes for the protein XAP5 circadian timekeeper. The *Arabidopsis thaliana* gene *At5g27430*, encoding the protein signal peptidase complex subunit 3B, shares 80.2% identity with the olive gene *OE9A057547*. We also reported a total of 18 candidate genes found in the different genomic regions corresponding to all significant SNPs found in one of the four GWAS analyses (Table S9). Their annotation and putative similarities correspond to 11 genes known in plant models and possibly to several transcripts (Table S9, Table S10). It is noticeable that the gene *OE9A037893* located on chromosome 15 encodes for a calcium-dependent protein kinase 4 (CPK4) whose putative function in potato is to regulate the production of Reactive Oxygen Species (ROS). These findings will provide a baseline for future candidate gene studies of FFD in olive.

## 4. Discussion

### Identification of three genetic clusters with varying flowering date in WOGBM

Consistently with previous studies (Diez et al., 2015; El Bakkali et al., 2019; Belaj et al., 2022), three genetic clusters were identified within the cultivated olive, based on the WOGBM. These clusters broadly correspond to the presumed geographical origins of the genotypes. The C1 group involved genotypes assumed to originate from the eastern Mediterranean, including Cyprus, Egypt, Greece, Lebanon and Syria. Group C2 consisted mainly of genotypes presumably originating from the central Mediterranean, encompassing Algeria, Croatia, France, Italy, Slovenia and Tunisia. The C3 group comprised genotypes putatively from the western Mediterranean, including Morocco, Spain and Portugal.

The comparison of genetic groups we obtained with the ones found in the same collection, WOGBM, but using SSR markers and another methodological approach (El Bakkali et al., 2019), and with the ones described in the WOGBC using either SSR (Diez et al., 2015) or EST-SNP markers (Belaj et al., 2022) revealed a general agreement in the composition of the groups (S1 File, Table S11, Table S12, Table S13). The concordance in terms of individuals assigned to each genetic group ranges from 66% to 85% for each respective group. The majority of individuals who were not assigned in our study were predominantly included in the non-assigned group from El Bakkali et al. (2019). The few discrepancies detected are assumed to result from differences in the approaches and markers employed. The STRUCTURE method (Pritchard et al., 2000) used by El Bakkali et al. (2019), Diez et al. (2015), and Belaj et al. (2022) relies on the assumptions of the absence of genetic drift, Hardy–Weinberg equilibrium, and linkage equilibrium between markers in ancestral populations (Pritchard et al., 2000), while the sNMF approach we used is not based on a genetic population model (Frichot et al., 2014). Moreover, the threshold of assignment to genetic clusters differs between the two methods. Even though these two methods usually converge (Frichot et al., 2014), it is not surprising that results may slightly differ.

Also, the markers used are possibly in different positions along the genome: SSR markers could be found in either coding or non-coding regions, while SNP markers in this study were selected to be located in coding regions or near them as we targeted annotated genes. Coding and non-coding regions are known to undergo different selection pressures (Jha et al., 2015). The two types of markers may have different evolution histories, with a higher mutation rate of SSRs compared to SNP markers (Fischer et al., 2017), that can result in different genetic structure signals. Moreover, our SNP data were not filtered for rare variants. Doing the analysis after applying a 5% MAF filter did not alter general structure, with more than 96% of similarities between the reported analysis and the one made after MAF filtration. Discordance was only due to some genotypes moving from a genetic cluster to the non-assigned group or vice versa (no shifts between genetic groups were observed) (Table S14, S1 File). This indicates that filtering for rare variants did not result in difficulty for classifying genotypes within one of the three genetic clusters.

Overall, in line with previous studies, we confirmed the existence of three distinct genetic clusters within cultivated olive. However, the boundaries between assigned and non-assigned genotypes are not fixed, as some genotypes assigned to a genetic cluster by a study could be found within the non-assigned in another one. Incorporating precise GPS coordinates of parent trees into our study could enrich our understanding of the genetic structure. Genotypes of the C3 group were closely related compared to C1 and C2 in the PCA plots. This finding aligns with the high level of genetic relatedness found between genotypes assigned to the Q1 cluster from Diez et al. (2015), representing western genotypes of MB.

A higher rate of non-assigned genotypes was observed in central MB compared to western MB and eastern MB. This suggests that admixture events may have occurred between genotypes from central MB and those from the western and eastern Mediterranean. Consistently with Diez et al. (2015), the non-assigned individuals were mainly from central and western MB.

### Marker-trait associations and potential candidate genes for flowering date

Distinct associated loci were detected in each of the four GWAS. Only three associated SNPs were consistent between at least two analyses. While a high value of heritability was estimated, these SNPs exhibited minor effects and accounted for a low proportion of the phenotypic variance. However, we must notice that the broad-sense heritability value was calculated based on a relatively small portion of the total variance of the trait, i.e. the part of variance explained by the genotypic effect only, as extracted from a mixed model, while the year and the interaction of genotype and year had high and significant effects. The combination of high heritability with few detected SNPs with low effects suggests that several other additional genomic regions could be involved in the genetic control of this trait.

Several factors may have prevented the detection of additional genomic regions. First, the genetic architecture of the studied trait is a key factor. A genetic architecture consisting of many loci with minor effects and/or rare variants with large effects can limit the power of GWAS to detect significant associations (Korte and Farlow, 2013). In our case, high accuracy values of genomic prediction were found with both RR- and LASSO-based models, even though the RR-based model performed significantly better than the LASSO-based model. This finding supports a polygenic genetic determinism underlying the flowering date trait in olive tree.

Second, the genomic data used can influence the detection power. Here, we used a capture sequencing approach, which targeted annotated genes rather than the Genotyping-by-Sequencing (GBS) method or whole-genome sequencing (WGS) which would have covered more exhaustively the genome, coding or non-coding. Given the high cost associated with WGS, the GBS method has been widely used as an alternative. While GBS offers a broader overview of the genome than capture sequencing, it often results in a high rate of missing data (Wang et al., 2020). This is due to the random digestion of the genome by restriction enzymes in GBS, leading to heterogeneous depth across genomic regions and variability in the coverage of loci between individuals (Elshire et al., 2011). In contrast, the capture sequencing approach used in the present work allowed to target identical genomic regions among individuals with high sequencing depth and limited missing data. Furthermore, capture sequencing of annotated genes enabled the identification of candidate genes after the GWAS, utilizing the annotation of associated loci. Even though WGS might be considered the best and most complete approach for GWAS studies, the capture sequencing chosen in this study appears to be an adequate compromise.

Third, the population size matters for the association detection power. A population size of less than 100 genotypes is usually considered too low to obtain a sufficient power of association detection (Hong and Park, 2012), even though the recommended population size depends on several factors, such as the genetic architecture of the trait with possible dominance and the extent of linkage disequilibrium (LD) (Hong and Park, 2012). The first association study in olive was performed using 96 olive genotypes sourced from the Turkish Olive GenBank Resources in Izmir, Turkey (Kaya et al., 2016). This study used a combination of SNP, AFLP, and SSR markers, totaling 1070 polymorphic loci, and focused on five traits related to yield. Subsequent GWAS studies, employing SNP data, have investigated the genetic determinism of various agronomic and morphological traits, making use of 183 genotypes (Kaya et al., 2019) or a large number of SNPs (428,320 SNPs) but 89 genotypes only (Bazakos et al., 2023). As our analysis benefited from a large dataset of 318 individuals genotyped with 118,948 SNPs, we can thus consider that those conditions are adequate to perform GWAS analysis.

Fourth, the power of detection depends on the frequency of SNP alleles within the studied population (Hong and Park, 2012). In WOGBM, the representation across Mediterranean regions of genotypes was unequal, with 25% of genotypes assumed to originate from Spain, 28% from Italy, and 18% from eastern MB only. This imbalance might result in a low frequency of alleles fixed in the eastern region in the whole population, even though they could be associated with the trait. It is noticeable that other types of populations, such as bi or multi-parental populations, although including less genetic diversity than collections, usually allow a better balance among allelic classes. Several studies based on bi-parental populations of apple tree have revealed a major QTL associated with flowering time that remained stable across populations (van Dyk et al., 2010) and was subsequently detected by GWAS (Watson et al., 2024). Therefore, combining investigations on bi-parental or multi-parental populations could complement the present study on WOGBM in the future. In this perspective, crosses between *Olivière* and *Arbequina* (Ben Sadok et al., 2013), have been created and could be used for such studies.

The analysis of linkage disequilibrium (LD) in the olive genome using SNP data from capture sequencing revealed a relatively rapid decay of LD. The average r2 value was relatively low (0.35), compared to the one reported using 57 olive cultivars sequenced via genotyping by sequencing technology (GBS) (Zhu et al., 2019). The LD decay distance observed in our study (∼100 bp) aligns closely with the one reported by Zhu et al., 2019 (∼85 bp) and is higher than that reported by D’Agostino et al., 2018 (∼25 pb), both studies using data from GBS. The LD decay of olive was relatively shorter than that found in pear (211 bp; Wu et al., 2018) and apple (161 bp; Duan et al., 2017). Considering the LD decay value in our study, the regions explored around the associated loci were extended. Three putative genes were localized in the explored regions. However, none of these genes has a known function related to flowering date, even though the *XCT* gene encodes functions related to the circadian clock and photomorphogenesis. Moreover, the gene found on chromosome 15 for a less robust association points towards a gene whose putative function is to regulate the production of Reactive Oxygen Species (ROS), known to be involved in dormancy release (Watson et al., 2024). These findings provide a baseline for future candidate gene studies of FFD in olive.

Another perspective of the present work would be to deepen the comprehension of the year effects and their interaction with genotypic effects on the FFD. Indeed, as previously found, flowering date is a highly heritable trait but also strongly depends on environmental conditions (Branchereau et al., 2023). Winter temperatures are particularly known to influence chilling fulfillment, which impacts FFD (Atkinson et al., 2013). Deciphering the genotype by year effects may lead to detect associations specific to a given year or environmental conditions, as previously demonstrated (Allard et al., 2016; Branchereau et al., 2023). As the WOGBM genotypes were phenotyped over seven years at the same experimental station (Tassaout, Morocco), testing associations for FFD per year will be interesting to assess environmental-specific associations. Additionally, phenotyping the same genotypes in various locations could be a longer-term perspective that would enhance differentiation between environments and facilitate the detection of environmental-specific associations and the exploration of FFD trait plasticity in response to environmental variations.

In conclusion, the BLUPs for the flowering date were associated with three loci only with minor effects, i.e. they accounted for a low proportion of the phenotypic variance. Considering the low effect and variance explained by the associated loci, these underlying genes should be approached with caution in the future. Altogether, our results suggest the implication of other genomic regions not being detected so far. The significantly higher accuracy of the RR-based model compared to the LASSO-based model in genomic prediction supports the hypothesis of a polygenicity of the trait. This knowledge could be further considered in olive breeding programs that will have to create new material combining optimal yield and flowering date adapted to future climatic conditions.

## 5. Materials and methods

### Plant materials

We used a panel of olive tree genotypes from the WOGBM. This collection is located at 31°49’10" N; 7°25’58" W (CRS: WGS84-EPSG:4326) in the Tassaout experimental station (Marrakech, Morocco), at an altitude of 465 meters above sea level (Abou-Saaid et al., 2022). The collection is initially composed of 554 accessions originating from 14 countries around the Mediterranean area. Characterization analyses using 20 SSR markers and 11 endocarp traits identified 331 unique cultivars within the collection (El Bakkali et al., 2019). The phenotyping was conducted on the 331 genotypes of the WOGBM collection, while genotypic data remained for 318 genotypes only after all data processing (see below).

### DNA extraction and genotyping

DNA was extracted from leaves using MATAB protocol and NucleoMag Plant Kit (Cormier et al., 2019). Libraries were constructed with NEBNext® Ultra™ II FS DNA Library Prep Kit (New England Biolabs, Ipswich, MA).

We constructed 333 individual genomic libraries from 330 accessions, thus including some experimental duplicates. Of the total sequenced samples three were duplicated from the same extraction and preparation, to assess the reproducibility of the experiment (S2 File, Table S15): *Leccino* (MAR0016), *Picual* (MAR00267), and *Picholine Marocaine* (MAR00540). These libraries were subject to capture experiments. We targeted the first 640 bp of each of the 55,595 annotated genes available by placing 1 to 4 probes (depending on gene length) of 80 bp each, with 0.5x tilling. The filtered set captured 16.8 Mb, including 210,367 baits representing 55,452 unique loci (Zunino et al., 2024). The Mybaits custom kits were designed and synthesized by Daicel Arbor Biosciences (Ann Arbor, Michigan, USA). Additionally, two genomic libraries, derived from the initial preparation of libraries but not subjected to the capture experiment, were sequenced: *Picholine* (MAR00196) and *Picholine Marocaine* (MAR00540), and were used as a control to estimate capture efficiency. All captured and non-captured libraries were pooled together in equimolar conditions. MGX-Montpellier GenomiX has performed the sequencing on an Illumina® NovaseqTM 181 6000 (Illumina Inc., San Diego, CA, USA). The detailed protocol was described by Zunino et al. (2024).

### SNP calling and filtering

We trimmed raw sequencing reads using FastP version 0.20.1 (Chen et al., 2018), where genotype *Aharoun* (MAR00447) was filtered out (quality reads below 30). The remaining reads were aligned to the reference genome of olive, Farga V2 (Julca et al., 2020), using the bwa-mem2 version 2.0 software (Vasimuddin et al., 2019). Duplicate reads were removed from sorted reads using picard-tools version 2.24.0. Alignments were then cleaned to keep only primary alignment, properly paired, and unique reads. The genotype *Azeradj Tamokra* (MAR00448) was removed due to its mapping rate of 0%. Finally, variants were called using the Genome Analysis Toolkit version 4.2.0.0 (Poplin et al., 2018) following GATK best practices. The final dataset comprises 64,835,479 variants across 333 samples. Data from the two non-captured libraries of *Picholine* (MAR00196) and *Picholine Marocaine* (MAR00540), were used to calculate the enrichment rate (the mean depth of targeted sequencing divided by the mean depth of non-captured sequencing). All the steps, from read cleaning to variant calling, were performed using the following Snakemake workflow: https://forgemia.inra.fr/gautier.sarah/ClimOlivMedCapture.

We removed the three biological replicates: *Unknown-VS2-545* (MAR00546 and MAR00547) and *Dhokar* (MAR00417), the three experimental duplicates: *Leccino* (MAR0016), *Picual* (MAR00267), and *Picholine Marocaine* (MAR00540), and the two non-enriched samples: *Picholine* (MAR00196) and *Picholine Marocaine* (MAR00540). This filter resulted in 325 genotypes being filtered to ensure data quality. We filtered out low-quality SNPs below a threshold of 200 and indels. We allowed a maximum of 3 SNPs within a 10 bp region and set the minimum mean depth per site at 8, with a maximum of 400. Additionally, the minimum mean depth per genotype was restricted to 8. We retained only biallelic SNPs. SNPs with a heterozygosity rate greater than 75% were removed. Loci with more than 10% missing data and samples with over 25% missing data were also excluded. Singleton SNPs were filtered out. The outcome dataset comprises 235,825 SNPs across 318 individuals. This set was used for genetic structure and PCA analyses. An additional filtration step consisting of setting a minor allele frequency filter of 0.05 was applied before the GWAS analysis, resulting in a set of 119,614 SNPs for the 318 individuals. The nuclear SNPs set comprises 119,600 variants (Table S2). This SNP set was used for the GWAS analysis, including a missing data imputation step followed by a minor allele frequency filter of 0.05 (see below).

### Phenotypic data and statistical analyses

Full flowering dates [Stage 65 according to the BBCH scale of olive tree (Sanz-Cortés et al., 2002)] have been recorded for the 331 genotypes of the WOGBM for seven years. Data from 2014 to 2019 were previously reported by Abou-Saaid et al., 2022. Additional data were collected in 2021 using the same methodology (Abou-Saaid et al., 2022). The collection exhibited varying numbers of repetitions per genotype, with each genotype being represented by a minimum of three trees. Some genotypes were represented by multiple trees because of synonymy and redundancy cases. For example, *Picholine Marocaine* was represented by 88 trees.

To account for the effect of years and possible interaction between years and genotypes on phenotypic data, three mixed models were tested and compared [see also (Abou-Saaid et al., 2022)]: (i) the model with the genotype as a random effect only; (ii) the model with the genotype as a random effect and the year as a fixed effect and (iii) the model with interaction “genotype × year” as a second random effect. The last model was the best model regarding the Akaike Information Criterion (AIC) (Akaike, 1974) and Bayesian Information Criterion (BIC) (Schwarz,1978) (Table S16, Table S17).

The equation of the best model is:

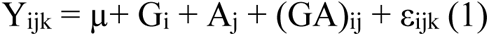

where Y_ijk_ represents the FFD value of tree k from genotype i in year j, µ denotes the overall mean of the population, G_i_ is the random effect of genotype i, A_j_ is the fixed effect of year j, (GA)_ij_ represents the random interaction between genotype i and year j, and ε_ijk_ represents the random residual error. the broad-sense heritability (H^2^) (Hühn et al., 1975) was estimated based on variance components:

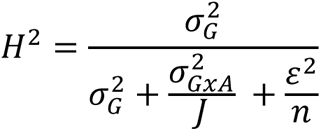

where 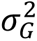 is the variance of genotype effect; 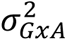 is the variance of interaction between genotype and year effect; ε^2^ is the variance of the residual term; J is the number of years and n is the mean number of observations per genotype and year.

The best linear unbiased predictor (BLUP) of the genotypic values of FFD for the 331 cultivars was extracted from the mixed model (1). The normality of BLUP of FFD genotypic values was tested using the Shapiro-Wilk test in R (Shapiro and Wilk, 1965).

### Population structure

To investigate the genetic structure of the cultivated olive collection under study, we used the dataset consisting of 235,825 SNPs from 318 genotypes. The genetic structure analysis was conducted using the sNMF approach (Frichot et al., 2014) implemented in the LEA R package (Frichot et al., 2015). This allowed us to estimate individual ancestry coefficients and determine the number of ancestral populations (K) within the dataset. We performed sNMF with K values ranging from 2 to 10. The smallest K value at which the cross-entropy did not significantly differ from that of K+1 was considered the most likely value of K.

Genotypes were assigned to genetic clusters based on their ancestry coefficients. If a genotype exhibited a minimum of 70% ancestry coefficient to a genetic cluster, it was assigned to that genetic cluster. Genotypes not reaching a 70% assignment to any of the genetic clusters are classified as non-assigned. To further investigate the genetic relationships among individuals, we performed a principal component analysis (PCA) to visualize their distribution within the population. The distribution of the genetic BLUP of FFD was compared between genetic groups using the Wilcoxon-Mann-Whitney test (Wilcoxon, 1945).

### Genome-wide association analyses

The association test was conducted between the BLUP of FFD genotypic values and the genomic data from the 318 genotypes of the WOGBM collection. The initial genomic dataset contained 119,600 filtered SNPs (Table S2), with 2.4% missing data. The missing data were imputed based on the genetic structure inferred by sNMF, using the LEA R package v3.11.3 (Frichot et al., 2015). The resulting imputed dataset was filtered for a minor allele frequency of 5%, resulting in 118,948 SNPs.

Three mixed models were tested and compared using the MM4LMM package (Laporte et al., 2022) to evaluate the inclusion of a random polygenic term and/or a fixed population structure effect in the model: i) the model with only polygenic effect (u), ii) the model with only genetic structure effect (Q), and iii) the model with both polygenic and genetic structure effects (u+Q). Two kinship matrices were tested for the covariance of the polygenic effect: the Weir and Goudet method (2017), implemented in the HIERFSTAT package in R (Goudet, 2005), and the VanRaden method (2008), implemented in the statgenGWAS package in R (Astle and Balding, 2009). VanRaden’s method is widely used in association studies, while Weir & Goudet is better suited to the structure of our dataset, especially considering the relatedness among certain genotypes (Goudet et al, 2018).

The most complete model equation was as follows:

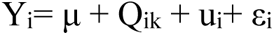

Where Y_i_ is the BLUP value for genotype i, Q_ik_ the fixed effect of the assignment of genotype i in structure group k, ui the random polygenic effect for genotype i and εi the random residual error. u_i_ ∼ N(0, 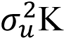), K being the genomic relationship (kinship). The best model was selected based on the Akaike Information Criterion (Akaike, 1974) and Bayesian Information Criterion (Schwarz,1978) (AIC and BIC; Table S6). The model that only included the random polygenic term was the best, regardless of the kinship matrix used to model its covariance, as it had the lowest values for both AIC and BIC. For further GWAS analysis, we thus used a model with the polygenic term only, but considering both the VanRaden, or Weir and Goudet methods for modeling the covariance of this polygenic effect.

The GWAS analysis was carried out using both single-locus and multi-locus models. For the single-locus model, we employed the MM4LMM package (Laporte et al., 2022), while for the multi-locus model, we utilized the MLMM approach, as proposed by Segura et al. (2012). MLMM is based on a forward and backward stepwise linear mixed model approach. In the forward steps, the most significant SNP detected in a step is incorporated into the model as a new cofactor before running again the GWAS, until reaching a defined threshold. Conversely, in the backward stepwise process, the least significant SNP from the list of candidates identified in the forward steps is removed from the cofactors at each step until only a single selected marker remains. The selected model was the one with the largest number of SNPs, which all have a P-value below the multiple-testing significance threshold as previously determined (Segura et al., 2012).

The combination of models (MM4LMM and MLMM) and kinships (VanRaden and Weir & Goudet) resulted in four distinct analyses. The significance threshold for MM4LMM was set at 5% false discovery rate (FDR). For MLMM, the threshold was established at 9.6 E-6, corresponding to the p-value of the least significant SNP in the initial run analysis of MM4LMM using the Weir & Goudet kinship matrix.

To calculate the variance explained by significant SNPs, likelihood-ratio-based R^2^_LR_ (Sun et al., 2010) was calculated for retained SNPs associated with the FFD trait.

### Looking for candidate genes

To include all SNPs in linkage disequilibrium (LD) in the region investigated for candidate genes, we estimated LD between SNPs using PopLDdecay V3.40 (Zhang et al., 2019) on a total of 235,825 SNPs from 318 genotypes (the same dataset used to study the genetic structure). The LD decayed at approximately 100 bp (r2 = 0.2). In order to encompass a larger genomic region, we extended the windows around the significantly associated SNPs by 1500 bases upstream and downstream of the SNP positions. We retrieved the list of genes within these defined intervals, along with their annotations and associated Gene Ontology (GO) terms reported by Julca et al. (2020), using the bedtools program v2.30.0 (Quinlan and Hall, 2010). Protein sequences of the genes found in these associated regions were further analyzed using BLAST against the UniProt database (The UniProt Consortium, 2023). Descriptions of these genes are provided in Table S10.

### Assessing accuracies of different Genomic Prediction models

We tested the accuracy of the genomic prediction of FFD BLUPs. For that, we used the same set of 118,948 SNPs of imputed data, previously used in the GWAS analysis, involving 318 individuals. Two genomic prediction models based on different regression algorithms to describe genetic architecture were tested. The ridge regression (RR) based model (Hoerl and Kennard, 1970), designed for scenarios with many minor effects, shrinks all marker effects toward 0 (but never truly 0) and the least absolute shrinkage and selection operator (LASSO) based model (Tibshirani, 1996), designed for scenarios with a limited number of major effects, enforces other effects to be exactly 0. The relative performance of RR or LASSO-based models could provide valuable information on the genetic architecture of the trait. Both models were implemented using the R/glmnet package (Friedman et al., 2010). Cross-validation to calibrate the shrinkage parameter λ was performed using a five folds cross-validation. Model accuracy was assessed by calculating the Pearson’s correlation between the observed values of the validation set (representing 1/5 of the total data) and the estimated values. One hundred iterations were conducted to estimate the distribution of model accuracy. The distribution of the accuracy values was compared between RR and LASSO-based models using the Wilcoxon-Mann-Whitney test (Wilcoxon, 1945).

## 6. Acknowledgments

We thank Hélène Vignes, Ronan Rivallan, and Anaïs Fossot for the laboratory help. Analyses were conducted on MESO@LR-Platform at the University of Montpellier with the help of members of the French Institute of Bioinformatics (IFB) - South Green Bioinformatics Platform. Laila Aqbouch PhD was founded by an IOC scholarship (N° 2021-03-PhD GRANT). This study was funded through Labex AGRO 2011 – LABX-002, project n° 2003-001 (under I-Site Muse framework) coordinated by Agropolis Foundation.

## 7. Contributions

Research Design: BK and AE. Laboratory experiment design: LZ and PM. Laboratory work: LA, FB, and PM. Resources, phenotypic data acquisition and curation: OA, AE and HZ. Genotypic data acquisition and curation: LA and GS. Data analysis and interpretation of results: LA, PC, GS, EC, BK, VS. PhD Supervision: EC, PC, GS, BK. Writing: LA, EC, PC. review & editing: All authors.

## 8. Data availability statement

Raw sequences data are available in the following database: ClimOliveMed; 2023;GenomiCOM: ClimOliveMed Genomic resources for research on adaptation of olive tree to climate change; European Nucleotide Archive; 2023-04-17; PRJEB61410. Scripts used in this study are available in the GitHub repository: https://github.com/laqbouch/Genetic_determinism_of_cultivated_olive.git

