## Supplementary information 1 for "Genome-wide association analysis of flowering date in a collection of cultivated olive tree"

### 1 UMR AGAP Institut, Univ Montpellier, CIRAD, INRAE, Institut Agro, Montpellier, France

### 2 Université Cadi Ayyad, Laboratoire Biotechnologie et Bio-ingénierie Moléculaire, FST Guéliz, Marrakech, Morocco

3 INRA, UR Amélioration des Plantes, Marrakech, Morocco

### 4 DIADE, Univ Montpellier, CIRAD, IRD, Montpellier, France

### 5 CIRAD, UMR AGAP Institut, F-34398 Montpellier, France.

### 6 INRA, UR Amélioration des Plantes et Conservation des Ressources Phytogénétiques, Meknès, Morocco

### 7 CBNMed, AGAP Institut, Montpellier, France

$These two authors contributed equally to this work

*These three last authors contributed equally to this work

**Supplementary figures**

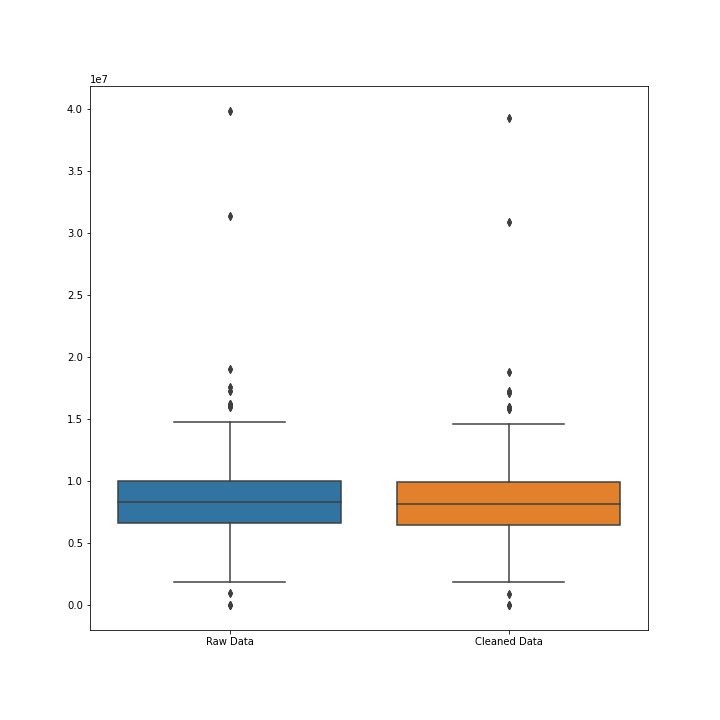

Figure S1. Boxplot of read counts of raw and cleaned data with a read quality threshold of 30 across the 333 sequenced libraries. Two genotypes, *Azeradj Tamokra* (MAR00448) and *Aharoun* (MAR00447), filtered before SNP calling were not included in the plot. The horizontal bar indicates the median value, and the boxplot the first and third quartile of each distribution, respectively.

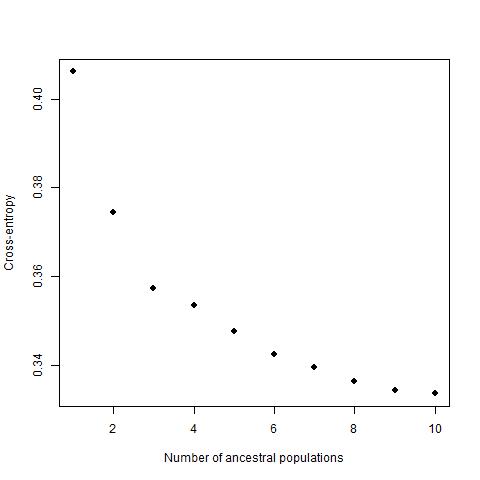

Figure S2. Cross-entropy criterion values by the number of ancestral populations from the sNMF Analysis (Frichot et al., 2014) for the 318 genotypes of Worldwide Olive Germplasm Banks of Marrakech (WOGBM) using 235,825 SNPs.

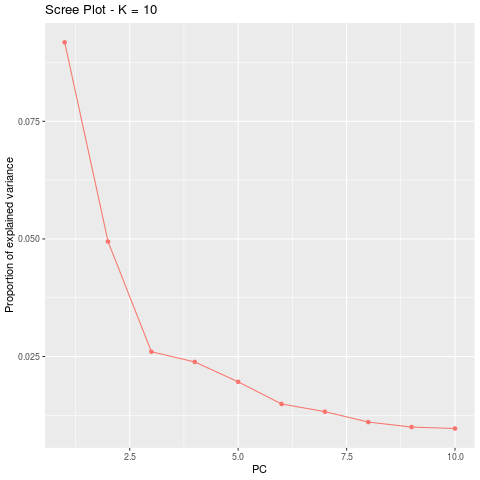

Figure S3. The proportion of genetic variability explained by the 10 first principal components in the principal component analysis (PCA).

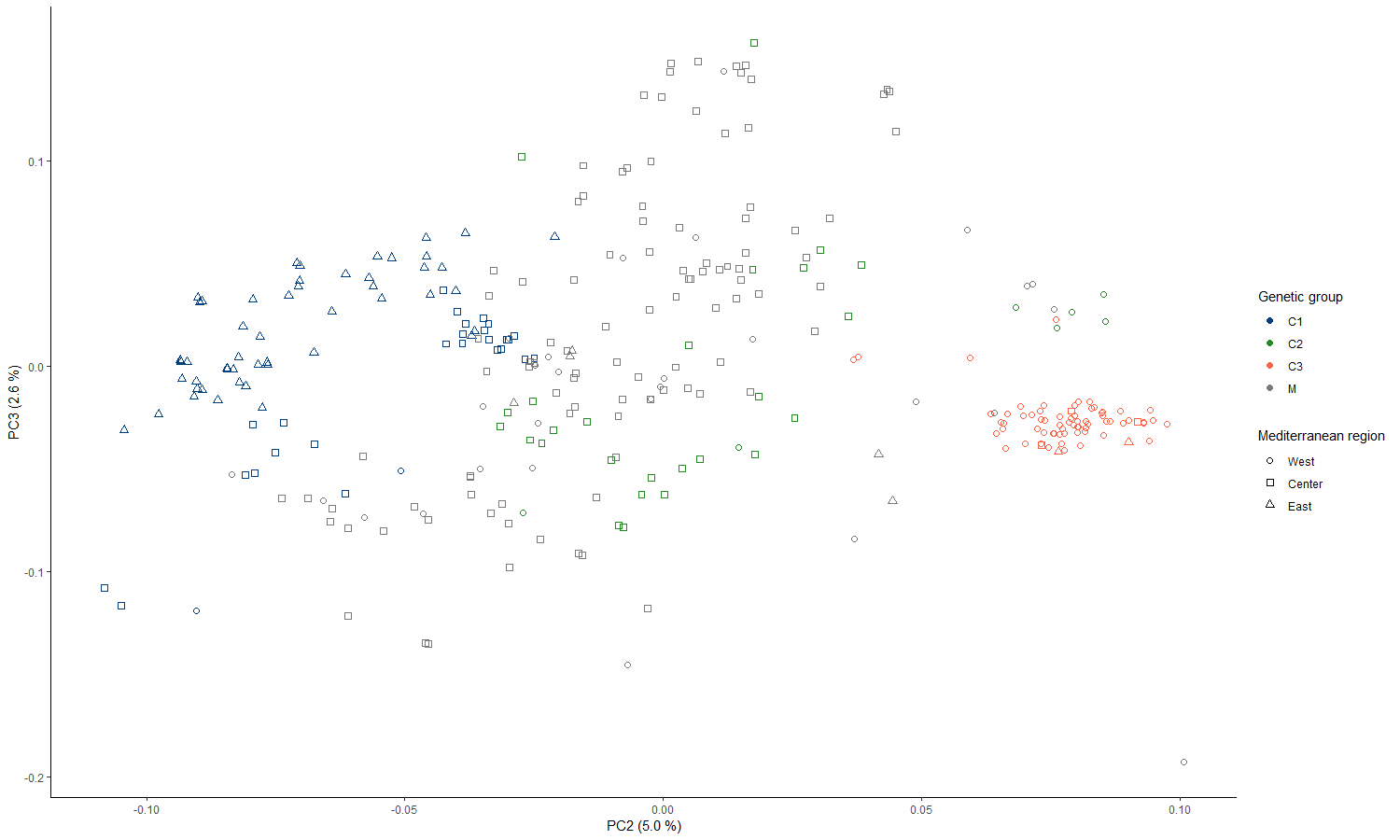

Figure S4. Projection of the 318 genotypes from the WOGBM on the second and third principal components of the PCA. Colors blue, green, and orange indicate the group to which each genotype was assigned (C1, C2, C3), and gray indicates the non-assigned genotypes (M). Circles, squares, and triangles indicate genotypes that are assumed to originate from the western, central, and eastern regions of the Mediterranean basin (MB), respectively. The east corresponds to Cyprus, Egypt, Greece, Lebanon, and Syria; the center corresponds to Algeria, Croatia, France, Italy, Slovenia, and Tunisia; and the west corresponds to Algeria, Croatia, France, Italy, Slovenia, and Tunisia.

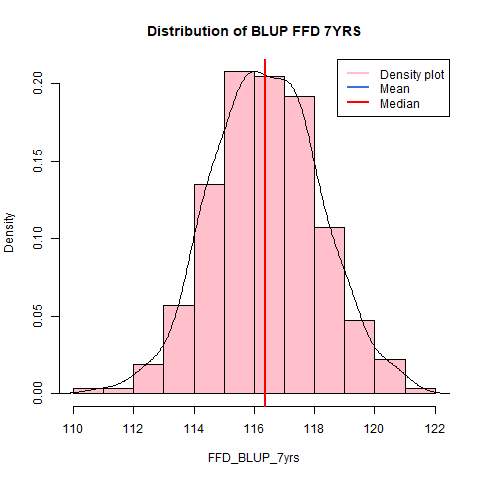

Figure S5. Density plot of the estimation of genotypic Best Linear Unbiased Predictors (BLUP) of full flowering date (FFD), estimated from a mixed linear model based on seven years of data, for 331 genotypes of the WOGBM collection. The blue vertical line indicates the mean, and the red one indicates the median. The continuous line indicates the distribution.

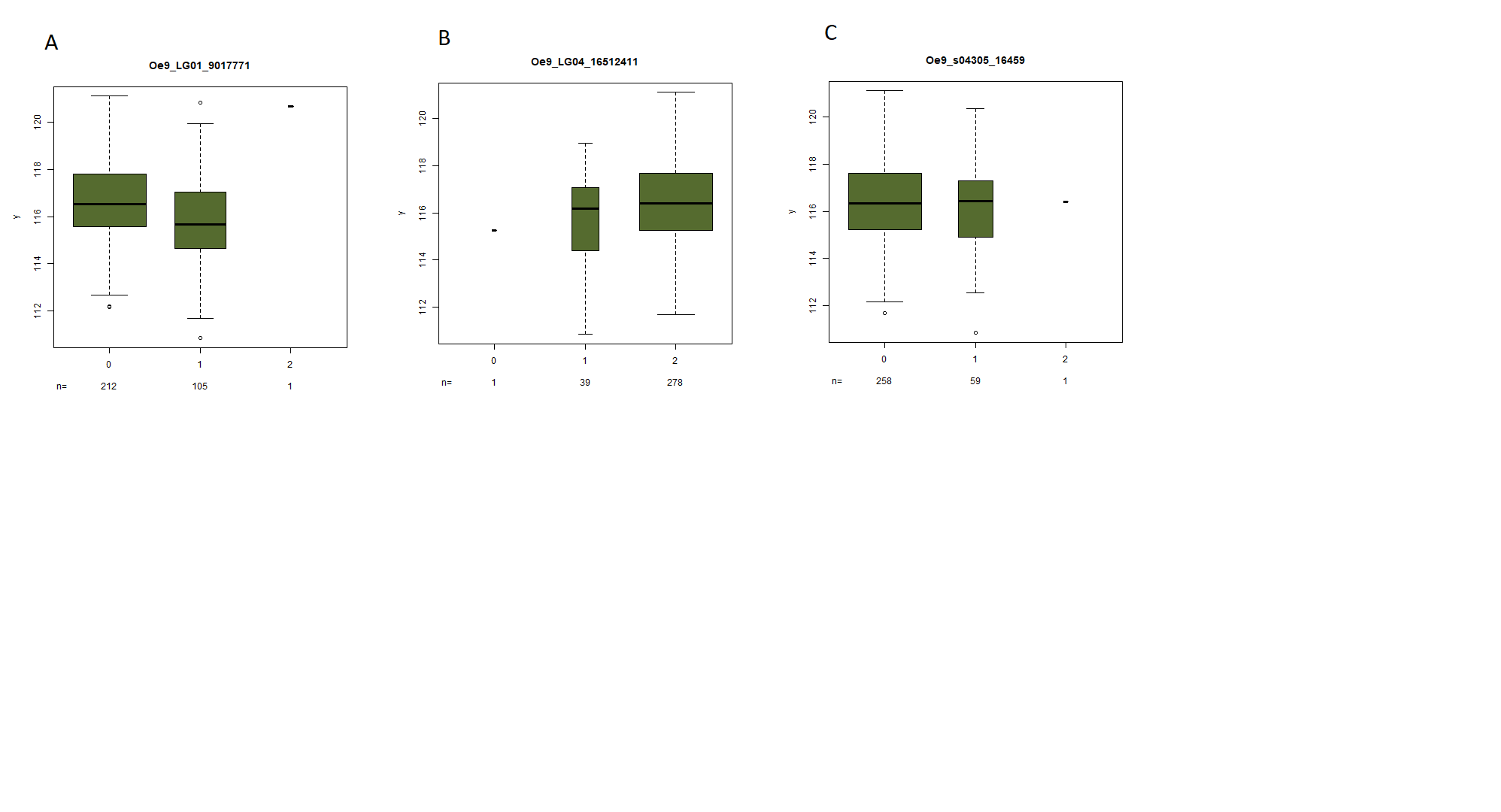
Figure S6. Boxplots of the genotypic BLUP values depending on the allelic classes (0 for the homozygotes for the reference allele, 1 for the heterozygotes, and 2 for the homozygotes for the alternative allele) for the three robust SNPs associated with genotypic Best Linear Unbiased Predictor (BLUP) of Full Flowering Date (FFD) identified by GWAS analysis. A. for *Oe9_LG01_9017771* SNP. B. *Oe9_LG04_16512411* SNP. C. *Oe9_s04305_16459* SNP. The number of individuals per class (n) is indicated below each graph.

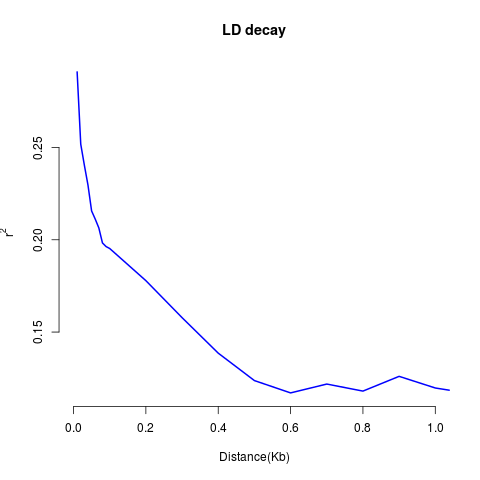

Figure S7. Decay of linkage disequilibrium (LD) in the WOGBM estimated by the correlation coefficient (r2) between consecutive SNP over a distance of 1000 bases, calculated with PopLDdecay V3.40 (Zhang et al., 2019).

**Supplementary tables**

Table S1. Mean depth values in captured and non-captured sequencing of the same genotypes, *Picholine* and *Picholine Marocaine*, with corresponding enrichment rate values. This table is in the supplementary xls file.

Table S2. Description of the successive steps of variant data filtering with the indication of the number of SNP and samples remaining after each filter, and the analyses performed on the last steps (estimation of the genetic structure, Genome-Wide Association Study and estimation of genomic-prediction-based models. This table is in the supplementary xls file.

Table S3. Estimated individual ancestry coefficients as defined by the sNMF Approach for K=3 (Frichot et al., 2014) for the 318 studied genotypes. Genotypes having more than 0.7 ancestry coefficients to a genetic cluster were assigned to it. Genetic group C1 included genotypes assigned to the K1 genetic cluster, C2 included genotypes assigned to K2, and C3 included genotypes assigned to K3, and non-assigned genotypes were denoted as M. This table is in the supplementary xls file.

Table S4. Percentage distribution of genotypes from different genetic groups (C1, C2, C3, and M) by their supposed geographical origin in three regions of the MB, information from Elbakhli et al. (2019). This table is in the supplementary xls file.

Table S5. Mean values of genetic BLUP of FFD per genetic groups (C1, C2, and C3) and pairwise comparisons based on the Wilcoxon-Mann-Whitney test (Wilcoxon, 1945). Levels of significance: ns (not significant); * (p<0.05); ** (p<0.01); *** (p<0.001). This table is in the supplementary xls file.

Table S6. Comparison of the Akaike Information Criterion (AIC) (Akaike, 1974) and the Bayesian Information Criterion (BIC) (Schwarz,1978) values of three linear mixed models that account for structure and/or kinship effects. The structure was considered as a fixed effect (as assessed by the ancestry matrix obtained from the sNMF run that exhibited the lowest cross-entropy value at the considered K, Q model), while the kinship was considered as the covariance matrix of a random effect separately (u model) or jointly (u+Q model). Two kinship matrices were considered Weir & Goudet (Weir and Goudet, 2017) and VanRaden Kinship (VanRaden, 2008). This table is in the supplementary xls file.

Table S7. Estimated values of genetic BLUP of FFD trait for the 318 studied genotypes. The name of each cultivar is indicated with its corresponding code in the WOGBM. This table is in the supplementary xls file.

Table S8. Results of GWAS analyses and characterization of SNPs Associated with genotypic BLUP of FFD trait using either the single-locus model MM4LMM or the multi-locus model MLMM: SNP name, chromosome or scaffold number, position in base pair, allelic composition (Ref indicates the allele of reference and ALT the alternative allele), minor allele frequency (MAF), model (MM4LMM or MLMM), kinship matrix used (Weir & Goudet or VanRaden), p-value for each SNP. This table is in the supplementary xls file.

Table S9. Annotation of genes found in the associated regions, corresponding to 1500bp upstream and downstream of each of the associated SNPs with genotypic BLUP of FFD: Chromosome or scaffold number; interval position of the associated region from the olive reference genome Farga V2 (Julca et al., 2020); Associated SNP position; candidate gene start and end positions, Transcript name; Gene ID; annotation and ontology term from the reference genome. This table is in the supplementary xls file.

Table S10. List of annotated genes found in the associated regions from the GWAS analysis and after a BLAST analysis using UniProt database (The UniProt Consortium, 2023): Transcript name, status (reviewed or not), protein name, gene name, organism on which the BLAST was successful, percent identity, e-value, functions. This table is in the supplementary xls file.

Table S11. The contingency table of the genotypes per genetic group obtained using SNP markers in the present study and using SSR markers from El Bakkali et al. (2019). C1, C2, C3, and M are the genetic groups considered in this study, while East, Center, West, and ssr_admixed are the genetic groups from El Bakkali et al. (2019). The general concordance was estimated by the percentage of genotypes attributed to a similar group in both studies. This table is in the supplementary xls file.

Table S12. The contingency table of the genotypes per genetic group obtained using SNP markers in the present study and using SSR markers from Diez et al. (2015). C1, C2, C3, and M are the genetic groups considered in the present study, while Q3, Q2, Q1, and Mosaic are the genetic groups from Diez et al. (2015). The general concordance was estimated by the percentage of genotypes attributed to a similar group in both studies. This table is in the supplementary xls file.

Table S13. The contingency table of the genotypes per genetic group obtained using SNP markers in the present study and EST-SNP markers from Belaj et al. (2022). C1, C2, C3, and M are the genetic groups considered in this study, while A, B, C, and EST_SNP_M were genetic groups from Belaj et al. (2022). The general concordance was estimated by the percentage of genotypes attributed to a similar group in both studies. This table is in the supplementary xls file.

Table S14. Contingency table of the genotypes per genetic structure obtained with SNP markers using data before and after filtering at 5% MAF. C1, C2, C3, and M are the genetic groups before filtering for the MAF. C’1, C’2, C’3, and M’ are genetic groups after filtering for the MAF. The general concordance was estimated by the percentage of genotypes attributed to a similar group before and after filtering. This table is in the supplementary xls file.

Table S15. Reproducibility of capture sequencing experiment using the percentage of different loci based on Variant Call Format (VCF) before and after variant filters for three genotypes: *Leccino*, *Picholine Marocaine*, and *Picual.* This table is in the supplementary xls file.

Table S16. Comparison of the three linear mixed models applied to the FFD phenotypic values collected on 331 genotypes and seven years. The three models account for: the genotype as a random effect only, the genotype as a random effect and the year as a fixed effect, and the interaction “genotype × year” as a second random effect. npar: number of parameters used in each model. AIC: Akaike Information Criterion (Akaike, 1974). BIC: Bayesian Information Criterion (Schwarz ,1978). logLik: The logarithm of the likelihood function. Chisq: Chi-square statistic. Df: Degrees of Freedom. Pr(>Chisq): p-value. Levels of significance: ns (not significant); * (p<0.05); ** (p<0.01); *** (p<0.001). This table is in the supplementary xls file.

Table S17. Analysis of variance of full flowering date values as a function of genotype, year, and “genotype x year” interaction. Df: Degrees of Freedom. Pr>F: p-value. Levels of significance: ns (not significant); * (p<0.05); ** (p<0.01); *** (p<0.001). This table is in the supplementary xls file.
