## Supplementary material for "Genome-wide association analysis of flowering date in a collection of cultivated olive tree": File_S1

### 1 UMR AGAP Institut, Univ Montpellier, CIRAD, INRAE, Institut Agro, Montpellier, France

### 2 Université Cadi Ayyad, Laboratoire Biotechnologie et Bio-ingénierie Moléculaire, FST Guéliz, Marrakech, Morocco

3 INRA, UR Amélioration des Plantes, Marrakech, Morocco

### 4 DIADE, Univ Montpellier, CIRAD, IRD, Montpellier, France

### 5 CIRAD, UMR AGAP Institut, F-34398 Montpellier, France.

### 6 INRA, UR Amélioration des Plantes et Conservation des Ressources Phytogénétiques, Meknès, Morocco

### 7 CBNMed, AGAP Institut, Montpellier, France

$These two authors contributed equally to this work

 *These three last authors contributed equally to this work

### S1 File: Supplementary discussion

**Comparison of genetic groups between studies of genetic structure**

We compared the genetic groups obtained from the genetic structure of our study with those obtained from SSR markers on the same collection, WOGBM, as presented in El Bakkali et al. (2019), as well as with SSR markers from Diez et al. (2015) and EST-SNPs from Belaj et al. (2020) on the WOGBC collection. We reported the numbers and the percentage of genotypes from our genetic groups within their respective genetic groups, based on common genotypes. Genetic groups were previously defined in the study of El Bakkali et al. (2019) and Diez et al. (2015), while for Belaj et al. (2020), only the estimated individual ancestry coefficients were published. To assign genotypes of this last study to genetic groups, we defined the same threshold as our study of 0.7.

The comparison between the genetic structure from our SNP data and the one obtained from SSR data by El Bakkali et al. (2019) resulted in a general concordance (68%) (i.e. the number of individuals matching their corresponding group divided by the total number of individuals) (Table S14) based on shared genotypes. Genotypes within the C1 group matched 60% of the eastern cluster identified by SSR markers. The remaining C1 genotypes were classified as part of the non-assigned group, except for *Karme* (MAR00640) and *Djlot_Tadmori* (MAR00617), which were found within the central group. Group C2 matched the central cluster, with 58% of genotypes. The remainder of C2 was classified as non-assigned by SSR. A difference was observed in the classification of the C2 and the non-assigned group (M) using SNP markers, compared to the central and the non-assigned group identified using SSR markers. The M group matches 62% of the non-assigned SSR group and 31% of the central group, while the rest of the genotypes are divided between the eastern and western clusters. The C3 group aligned to the western cluster, with 93% of genotypes matching between the two groups. The other 7% were classified as table non-assigned by SSR markers.

A general concordance (66%) was also found between the genetic groups of our study and the ones obtained using SSR markers from Diez et al. (2015) on WOGBC (Table S15) based on 73 shared genotypes. The C1 and C3 genotypes aligned with Q3 and Q1, with 69% and 89% of genotypes matching, respectively, while the remaining genotypes were within the SSR mosaic group (equivalent to our M group). Only two genotypes were categorized as part of the C2 group based on SNP markers, while they were classified as Mosaic using SSR markers. The genotypes in the M group were shared by both the Q2 and the Mosaic group, with only one genotype belonging to the Q1 group.

A general agreement (85%) was found between the genetic groups of our study and the ones obtained from EST-SNP markers by Belaj et al. (2022) on the WOGBC (Table S16) based on 99 shared genotypes. We used the same threshold as our study of 0.7 to assign genotypes to genetic clusters. The C1 and C3 genotypes matched the A and C groups of EST-SNP, with 73% and 98% of concordance, respectively. The remaining genotypes in these groups were classified as non-assigned by EST-SNP. Among the four genotypes in our C2 group, only one was identified as part of the B group by EST-SNP. The M group shared 79% of genotypes with the non-assigned group identified by EST-SNP, while the remaining genotypes were assigned to either the B or C groups.

**Comparison of genetic groups before and after filtering for minor alle frequency**

We compared the genetic groups obtained from the genetic structure of our data before and after applying a 5% minor allele frequency (MAF) filter. To assign genotypes after MAF filter, we defined the same threshold as our study of 0.7.

A general agreement (96%) was found between the genetic groups of our study based on data before and after MAF filtering. The C1, C2, and C3 group of genotypes from data before MAF filtering matched the C’1, C’2, C’3 group of genotypes from data after MAF filtering, with 99%, 91%, and 99% of concordance, respectively. The remaining genotypes were classified as non-assigned (M’). The M group shares 93% of genotypes with M’ group, while the remaining genotypes were assigned to either the C’1, C’2, or C’3 groups.
