## Supplementary material for "Genome-wide association analysis of flowering date in a collection of cultivated olive tree": File_S2

**S2 File. Supplementary methods**

**Assessment of the reproducibility of capture sequencing experiment**

To assess the reproducibility of the capture sequencing experiment, we replicated three genotypes: *Leccino* (MAR0016), *Picual* (MAR00267), and *Picholine Marocaine* (MAR00540). We calculated the percentage of differences between replicates by dividing the number of different loci by the total number of tested loci, excluding missing alleles. This is done on raw VCF after variant calling (64,835,479 variants) and for the filtered VCF (we reproduced the same VCF filtering pipeline as described before (Table S2), except for the minor allele frequency and non-nuclear data). The error rate of the experiment is approximately 5.96% when using the raw VCF data, but it decreases to 2.5% when using the filtered VCF data (see Table S17).
